# Novel insights into the genetic relationship between growth and disease resistance in Pacific salmon

**DOI:** 10.1101/455196

**Authors:** Agustin Barría, Andrea. B. Doeschl-Wilson, Jean P. Lhorente, Ross. D. Houston, José. M. Yáñez

## Abstract

**Background:** Breeding for disease resistance has become a highly desirable strategy for mitigating infectious disease problems in aquaculture. However, knowledge of the genetic relationship between resistance and other economically important traits, such as growth, is important to assess prior to including disease resistance into the breeding goal. Our study assessed the genetic correlations between growth and survival traits in a large bacterial infection challenge experiment. A population of 2,606 coho salmon individuals from 107 full-sibling families were challenged with the bacteria *Piscirickettsia salmonis*. Growth was measured as average daily gain prior (ADG0) and during (ADGi) the experimental infection and as harvest weight (HW). Resistance was measured as Survival time (ST) and binary survival (BS). Furthermore, individual measures of bacterial load (BL) were assessed as new resistance phenotypes and to provide an indication of genetic variation in tolerance in salmonid species.

**Results:** Significant moderate heritabilities were estimated for ADG0 (0.30 ± 0.05), HW (0.38 ± 0.03), and for the survival traits ST (0.16 ± 0.03) and BS (0.18 ± 0.03). In contrast, heritabilities for ADGi and log-transformed BL were low (0.07 ± 0.02 (significant) and 0.04 ± 0.03, respectively), although these increased to moderate significant levels (0.20 ± 0.09 and 0.12 ± 0.05, respectively) when traits were assessed in survivors only. Significant and favorable genetic correlations were found between ADG0 and the growth traits ADGi (0.40 ± 0.16) and HW (0.64 ± 0.09), as well as with resistance as ST (0.43 ± 0.18), indicating that fish with higher genetic growth rate early on and prior to infection not only tend to maintain their genetic growth advantage until harvest, but also tend to grow faster and survive longer during infection. Furthermore, no robust unfavorable genetic correlations between ADG0 and any of the other traits considered in this study, in particular BL, was identified. Adding log BL as covariates into the models for growth under infection and survival provided an indication for genetic variation in tolerance.

**Conclusions:** These results suggest that selective breeding for early growth would be expected to simultaneously increase survival time and growth performance during an infection with *Piscirickettsia salmonis* after accounting for variation in bacterial load, and harvest weight in this coho salmon population, without negatively impacting on pathogen burden.

## Background

Chile is the main producer of farmed coho salmon (*Oncorhynchus kisutch*) globally, with approximately 90 % of the total production [1]. In common with other intensive production systems, the health status of farmed coho salmon is a major concern for profitability and animal welfare. One of the main sanitary issues affecting the Chilean coho salmon industry is Salmon Rickettsial Syndrome (SRS), caused by *Piscirickettsia salmonis*, a facultative intracellular bacteria which was first isolated in Chile [2]. SRS is responsible for great economic losses, either directly through mortality, or indirectly through treatment costs or reduction on fish performance. During the first half of 2016, *P. salmonis* was responsible for 53 % of mortalities attributed to infectious disease in farmed coho salmon [3]. Current strategies to control SRS, such as vaccines and antibiotics, have not been fully effective in tackling this disease under field conditions [4].

Selective breeding for improved resistance to infectious diseases is a feasible and potentially more sustainable strategy for the long-term control of disease outbreaks in both livestock and aquaculture species [5]. To date, salmonid breeding programs typically use disease challenge testing of relatives of the selection candidates to enable breeding values estimations and genetic improvement of disease resistance [6,7]. Accordingly, there is an increasing number of studies aimed at quantifying and dissecting the host genetic variation for disease resistance by measuring survival after exposure to diverse infectious pathogens [6,8–10]. Previous studies have demonstrated significant genetic variation for resistance to *Piscirickettsia salmonis* in Atlantic salmon (*Salmo salar*), rainbow trout (*Oncorhynchus mykiss*) and coho salmon using disease challenge data, with heritability estimates ranging from 0.11 to 0.62 [11–16].

From an economic perspective, growth, SRS resistance and flesh color are key traits to be included into the breeding goal of Chilean salmon [17]. Before including all these traits simultaneously into the breeding objective, knowledge of the genetic correlations between traits is needed. A positively correlated response to selection between growth at harvest and flesh color has been reported in a Chilean coho salmon breeding population [18]. Moreover, although [15] estimated a genetic correlation not significantly different from zero between SRS resistance (defined as day of death) and body weight in Atlantic salmon, the same authors reported a negative genetic correlation of −0.50 ± 0.13 between SRS resistance (defined as day of death) and harvest weight in coho salmon [12]. However, little is known about the relationship between growth prior to and during infection and SRS resistance.

In aquaculture, disease resistance is commonly defined using host survival data (measured as binary or day of death) after being exposed to an infection, either in an experimental challenge [6,12,14,19,20] or under field conditions [10]. However, this definition captures two different host response mechanisms to infections under potential genetic regulation, i.e. (i) the ability of the host to restrict pathogen invasion or replication (best described by within-host pathogen load) and (ii) the ability of an infected host (with a given pathogen load) to survive the infection [21]. Studies that include both types of mechanisms often refer to the first trait as ‘resistance’ and to the second trait as ‘tolerance or endurance’ [22–27]. Clearly, resistance and tolerance /endurance contribute both positively to an individual’s ability to survive an infection. However, at the population level, tolerant individuals that tend to harbor and can cope with high pathogen load are undesirable, as these would be expected to shed more infectious material and thus to cause a higher disease threat to individuals in the same contact group [28]. Thus, in order to minimize disease prevalence and mortality due to disease in a population, a fuller understanding of the relationship between within-host pathogen load and mortality is required. However, measures of individual pathogen load are rarely available in large scale genetic studies, especially in aquaculture. In the present study we aimed to overcome this shortcoming by quantifying bacterial load as a measurement of disease resistance in coho salmon families with extremes survival rates after infection with *P. salmonis*. We then evaluated the phenotypic association and genetic correlation of this trait with SRS survival traits and growth rate.

The aim of the current study was to provide a better understanding of the genetic relationship between growth and SRS resistance traits in coho salmon. Specifically, the objectives were (i) to quantify the level of genetic variation for growth both prior to and during an infection with *P. salmonis*, and for harvest weight, in a coho salmon breeding population, (ii) to assess the genetic correlations between these growth traits and survival traits in a large SRS infection challenge experiment, and (iii) quantify the level of genetic variation in individual bacterial loads as an alternative phenotype for disease resistance, and their relationship with survival and growth traits.

## Methods

### Breeding Population

The study was performed on a coho salmon (*Oncorhynchus kisutch*) population from the 2012 year-class, belonging to a genetic improvement program established in 1997. This breeding program is owned by Pesquera Antares and is managed by Aquainnovo (Puerto Montt, Chile). The breeding population consists of two sub-populations, depending on the spawning year. Both sub-populations have been selected for eight generations for harvest weight (HW) using best linear unbiased prediction (BLUP). Summary statistics for age at tagging (AT), weight at tagging (WT) and HW for population with spawning at even years are described in supplementary material (Table S1). More details about this breeding population are described in [12] and in [18].

### Experimental Challenge

The study comprised a total of 2606 individuals belonging to 107 maternal full-sib families (60 half-sib families) from the 2012 spawning year from the breeding nucleus described above. Prior to challenge, each fish was marked with a Passive Integrated Transponder (PIT-tag), placed on the abdominal cavity for genealogy traceability during challenge test. Body weight was measured for each fish prior to PIT-tag placement. The average weight at tagging was 5.5 g (SD = 1.02 g) and the mean age at tagging was 218 days (SD = 3). After tagging, fish were kept in communal tanks with fresh water at 13°C and then transferred to salt water [31 ppt] with an initial density of 15Kg/m^3^.

The strain of *P. salmonis* used in the current study was isolated in 2012 and purchased from ADL Diagnostic Chile Ltda, (Puerto Montt, Chile). A lethal dose 50 (LD50) was calculated from an independent assay prior to the main experimental challenge. For this pre-challenge, a total of 80 fish were randomly selected from the population. Four different dilutions (1/10, 1/100, 1/1000 and 1/10000) of the *P. salmonis* inoculum were evaluated on twenty individuals per dilution. Fish were intraperitoneally (IP) injected with a total volume of 0.2 ml / fish. Mortality was recorded daily. This preliminary assay lasted 26 days. A LD_50_ of 1:680 was estimated as described in [29] for use in the main challenge,

For the main challenge test, fish not used for the pre-challenge were distributed amongst three tanks (7 m^3^) with salt water concentration of 31 ppt. A mean of eight individuals (ranging from 1 to 18) per family were allocated into each tank. Each challenged family had on average 25 individuals (from 11 to 45), which were distributed across all three tanks. After an acclimation period of 12 days, fish were anesthetized using 30 ppm of benzocaine and body weight was measured for each individual. The mean initial body weight (IW) at the inoculation procedure was 279 g (SD = 138 g). The infection of each fish was then performed through an IP injection of 0.2 ml / fish of the LD_50_ inoculum. To ensure the absence of any other pathogens in the population, a sample of 30 fish from the full-sib families were randomly tested. Quantitative real-time PCR (qRT-PCR) was used to test for and exclude the presence of Infectious Salmon Anaemia Virus (ISAV) and Infectious Pancreatic Necrosis Virus (IPNV). Conventional PCR and culture was used to test for and exclude the presence of *Flavobacterium psychrophilum*, and an immunoflouresence antibody test (IFAT) was used to test for and exclude the presence of *Renibacterium salmoninarum*.

Mortalities returned to approximately baseline levels at day 47 which was maintained for three days. Thus, the experimental challenge continued for 50 days post injection. Mortalities were removed from each tank daily. At the end of the challenge test, all surviving fish were euthanized. Head kidney samples were taken from all dead and surviving fish and storage in RNALater at 4°C for 24 h and then transferred to - 80°C for longer term storage. Body weight at the day of death (FW) was measured for each individual. The disease challenge and measurements were all performed on Aquainnovo’s Research Station, in Lenca River, Xth Region, Chile.

A necropsy assay and qRT-PCR were performed on all dead fish, to confirm the presence of *P. salmonis* as the likely cause of death. The presence of *Vibrio ordalli*, *Renibacterium salmoninarum* and IPNV were discarded as mentioned above. Summary statistics for age and weight at tagging (AT, WT) and initial and final weights (IW and FW) for the challenged population is shown in Table S2.

### Growth and mortality phenotypes

The survival time (ST), measured as the day of death after IP injection was recorded for each fish. Values ranged from 1 to 49 depending on the time of the event. Individuals that survived until the end of the challenge were assigned a ST value of 50. Binary survival (BS) was defined as 1 if the fish survived until the end of the challenge or 0 if the fish died during the experimental challenge.

Growth rate was measured prior to and during *P. salmonis* infection. Average daily gain (g/day) before experimental challenge (ADG0) was measured as the difference between body weight at the time of IP injection and body weight at the time of tagging divided by the number of days (ranging from 239 to 251 days, depending on the family) between both procedures. Average daily gain during infection (ADGi) was calculated by the difference of weight at the day of death (or termination of challenge in the case of survivors) and weight at IP injection divided by the number of days between both timepoints. In addition to the records from the challenged individuals, harvest weight (HW) records from 41,597 fish with known relatedness to the challenge individuals (established through pedigree records) were available and included in the genetic analyses.

### Quantification of bacterial load

A subset of 740 individuals from 33 full-sib families were selected from the population for measurement of bacterial load. Using data from the experimental challenge, these families were selected based on their estimated breeding values for ST using *best linear unbiased prediction* (BLUP). More specifically, the 16 and 17 families with the highest and lowest estimated breeding values, respectively, were chosen and are hereafter referred to as the most and least resistant families, respectively. All 33 families were equally represented in each tank. Bacterial load (BL) from these individuals was quantified through amplification of the 16S gene by qRT-PCR. The gene copy number was log-transformed and corrected by the six copies of 16S in *P. salmonis* [30]. The log-transformed BL (LogBL) was then used as a measure of host resistance to *P. salmonis* infection [25,31].

### Statistical Analysis

Bivariate linear animal models were used to estimate the variance and covariance components among the traits. Bivariate models were defined as follows:

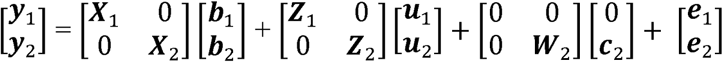

Where, **y_1_** and **y_2_** are vectors of phenotypic records measures in animals for either ST, BS, LogBL, ADG0, ADGi and HW; **b_i_** are vectors of fixed effects, for HW the contemporary group of sex:cage:year was included as factor, and harvest age, with marking age (MA) and marking weight (MW) fitted as covariates. For all remaining traits, sex and tank were included as fixed effects, and MW and MA were fitted as covariates. To account for different recording times for body weight and bacterial load in survivors and non-survivors, binary survivor / non-survivor status at the end of the challenge was included as fixed effect for ST, ADGi and LogBL. The variable ADG0 was included as covariate in the bivariate models between ADGi, logBL, ST and BS that did not also include ADG0 as response variable. Similarly, the effect of LogBL was included for ADGi, ST and BS that did not include LogBL as response variable; **u_i_** and **e_i_** are vectors of random animal genetic and residual effects, respectively, **c_2_** is the vector of random environmental effect associated with common rearing of full-sib families for HW prior to tagging; **X_i_** and **Z_i_** are the design matrices for the corresponding fixed and random effects for both traits, and **W_2_** is the design matrix for HW.

The random effects associated to each animal and residual effects, in conjunction with common environment effect for HW, were assumed to be normally distributed according to:

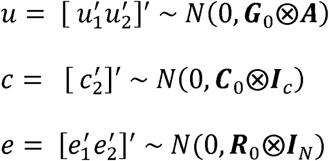

Where, **A** is the matrix of additive genetic kinship among all the fish included in the pedigree; **I**_C_ and **I**_N_ are the identity matrix of dimension **C** and **N**; ⊗ indicates the direct operator of the products; the matrices **G**_0_ and **R**_0_ denote the variances and co-variances of 2 x 2 additive genetic and residual effects, respectively. Common environment effect was evaluated for each trait using a single-trait likelihood ratio test [32]. This effect was significant only for HW (P < 0.05), and therefore included in the bivariate models when HW was analyzed. Thus, **C**_0_ represents a 1 x 1 scalar of common environment effect for HW. Given that HW was recorded on a different population of individuals to the challenge population, environmental covariance was set to zero in the **R**_0_ matrix. The parameters in the bivariate mixed linear models were estimated using the restricted maximum likelihood method (REML) implemented in ASREML version 3.0 [33].

To assess the influence of different recording times for body weight and bacterial load between survivors and non-survivors on (co-)variance estimates, bivariate analyses for logBL, ADG0, ADGi, and HW were repeated for subsets of data containing either survivors or non-survivors only. Similarly, bivariate analyses for ST were also performed exclusively for non-survivors.

### Heritability and genetic correlations

The following formula was used to estimate heritability values for the different traits:

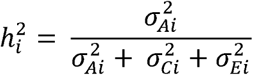

Where, i is either ST, BS LogBL, ADG0, ADGi or HW, 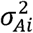 are the additive genetic variance of matrix **G**_0_, 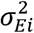 are the residual variances from the matrix **R**_0_ and 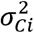 is the common environmental effect associated with the full-sib families (only for HW).

The genetic correlations (**r**_xy_) among traits were calculated as follows, according to [34]:

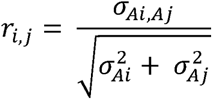

where, *σ_Ai,Aj_* corresponds to the additive genetic variance between the traits evaluated in the bivariate model, 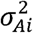 corresponds to the additive genetic variance of trait *i* and 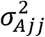 corresponds to the additive genetic variance of trait *j*.

## Results

### SRS experimental challenge

Typical clinical signs and pathological lesions associated with a *P. salmonis* infection were observed after IP injection. These signs included inappetence, lethargy and pale gills [4]. Mortality began on day 10 post IP injection. Dead individuals showed a swollen kidney, splenomegaly and yellowish liver tone (typical symptoms of SRS infection; [4]). During the 50 days of challenge, the three replicate tanks reached a cumulative mortality of 35.6, 42.5 and 37.7%, respectively.

Figure 1A shows the observed mortality from all 107 challenged families, ranging from 5.0 to 81.8 %, with an average mortality of 38.5 %. The most susceptible / resistant families selected for bacterial quantification are highlighted, with a mean mortality of 63 and 17%, respectively. The proportion of survivors and non-survivors among these 33 extreme families are shown in Figure 1B. Except for one family, all families contained both survivors and non-survivors, although the percentage of survivors was considerably higher in the most resistant families (ranging from 67 to 100 % compared to 19 to 48 % in the most susceptible families).

**Fig. 1.**
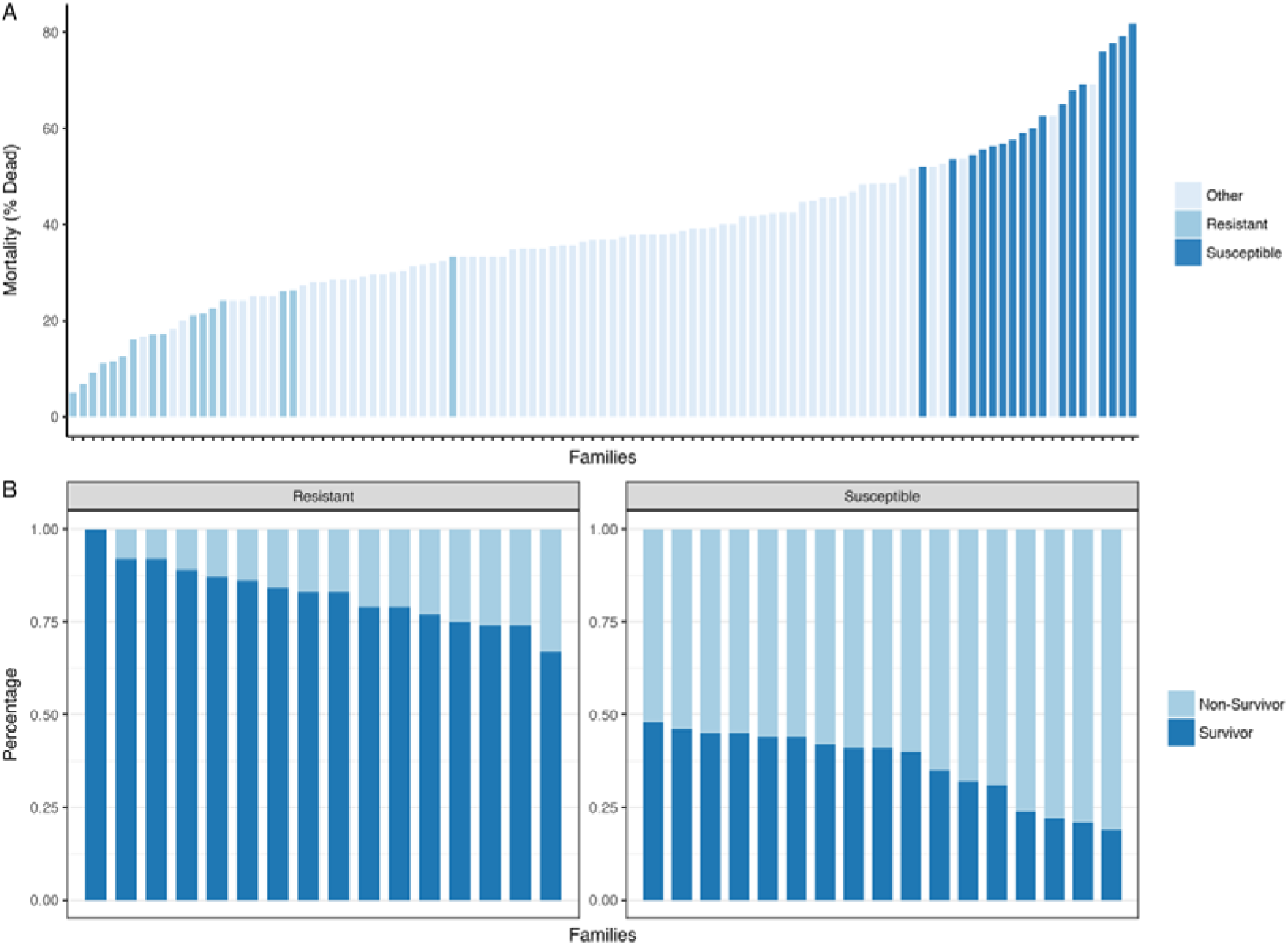
Observed mortalities for all the 107 coho salmon families challenged with *P. salmonis*. The 16 most resistant and 17 most susceptible families for which bacterial load was quantified are highlighted in light and dark grey, respectively (Fig 1A). Percentage of survivors and non-survivors for each of the 16 most resistant and 17 most susceptible families selected after experimental challenge against *P. salmonis* (Fig. 1B).

### Phenotypic variation for resistance and growth traits

Table 1 shows summary statistics of the phenotypic variation observed for the different measured traits. Prior to the challenge test, individuals gained on average 1.11 g/day (SD = 0.56) in body weight, ranging between 0.34 and 2.62 g/day. However, this average daily growth rate was reduced by almost half (0.69 g/day ± 1.65), and had far higher phenotypic variation during the infection process. Some individuals continued to grow (maximal gain of 6.36 g/day), whereas others experienced a weight loss (maximal weight loss of 5.85 g/day). The distribution for ST was right skewed, with 61% of fish having a ST value of 50 (Figure S1), i.e. they survived the experimental challenge. Amongst non-survivors, fish survived on average 41 days (SD=10) after IP injection, with a minimum value of 10 days and maximum value of 49 days.

**Table 1.** Summary statistics for average daily gain prior P. salmonis challenge (ADG0), during challenge test (ADGi), harvest weight (HW), log10 of bacterial load (LogBL), day of death (ST) and binary survival (BS) in the coho salmon (Oncorhynchus kisutch) breeding population used in the present study.

| Trait | Mean | SD<sup>a</sup> | CV<sup>b</sup> (%) | Min<sup>c</sup> | Max<sup>d</sup> | Number of records |
| --- | --- | --- | --- | --- | --- | --- |
| ADG0 (g/day) | 1.11 | 0.56 | 50.29 | 0.34 | 2.62 | 2605 |
| ADGi (g/day) | 0.69 | 1.65 | 239.13 | -5.85 | 6.36 | 2597 |
| HW (Kg) | 6.37 | 1.32 | 20.72 | 0.05 | 7.5 | 41597 |
| LogBL | 1.44 | 0.51 | 35.42 | 0.0 | 2.36 | 740 |
| ST | 41.56 | 10.01 | 24.09 | 10 | 50 | 2606 |
| BS | 0.61 | 0.49 | 80.33 | 0.0 | 1.0 | 2606 |

<sup>a</sup> Standard deviation
<sup>b</sup> Coefficient of variation
<sup>c</sup> Minimum
<sup>d</sup> Maximum

Bacterial load quantification of 740 fish from 33 families showed that 6.5 % (n = 48) of individuals had a *P. salmonis* load below the qRT-PCR detection threshold; these fish were considered as bacterial-free. All these individuals survived the experimental challenge and belonged to the resistant families. The bacterial load measured on the log_10_ scale in the other 692 animals ranged from 0.81 to 2.36 with an average load of 1.53 log units (SD = 0.33). From these 692 individuals, 55 % were survivors, while the rest of the animals died during the experimental challenge. Harvest weight,obtained from 41,597 commercial fish with linked pedigree to the challenged population, had a mean of 6.36 kg (ranging from 0.05 to 7.5 kg).

### Comparison of traits measured in survivors and non-survivors

Binary survival (BS), added as fixed effect in the statistical models, had a significant effect on ADGi and logBL (p < 0.001) (Table 2). Survivors grew on average by 2.09 ± 0.56 g/day faster than non-survivors during the experimental challenge and also experienced on average a 0.64 ± 0.03 times lower bacterial load (in log units) (data not shown). Furthermore, genetic correlations between the traits ADGi, and LogBL measured in survivors and non-survivors, respectively, were low (0.026 ± 0.64 for ADGi and 0.001 ± 0.39 for logBL), indicating that these traits may be considered as genetically different traits in survivors and non-survivors. Interestingly, the same was true for growth prior to infection, where the genetic correlation between ADG0 in survivors and non-survivors was 0.032 ± 0.21 (Table S3).

**Table 2.** Summary of the variables used for each mixed models and the significance effect

| Trait | Pooled Animals |  |  |  |  |  |  |  |  |
| --- | --- | --- | --- | --- | --- | --- | --- | --- | --- |
|  | Sex | Tank | MA<sup>a</sup> | MW<sup>b</sup> | ADG0<sup>c</sup> | LogBL<sup>d</sup> | BS<sup>e</sup> | CG<sup>f</sup> | HA<sup>g</sup> |
| ADG0 | ** | NS<sup>k</sup> | -2.2E-02(5.5E-03)** | 1.4E-01(1.1E-02)** | NA<sup>l</sup> | NA | NA | NA | NA |
| ADGi<sup>h</sup> | ** | ** | 3.4E-03(1.0E-02)NS | -2.9E-02(2.8E-02)* | 8.8E-02(5.1E-02)** | -7.7E-02(4.5E-02)NS | 2.1(5.5E-02)** | NA | NA |
| HW<sup>i</sup> | NA | NA | -2.1E-02(4.1E-03)** | 92.8(7.4)NS | NA | NA | NA | ** | 1.4E-02(2.8E-03)** |
| LogBL | ** | ** | -7.3E-04(5.3E-03)NS | 2.1E-02(1.5E-02)* | -3.8E-02(2.7E-02)** | NA | -6.4E-01(3.0E-02)** | NA | NA |
| ST<sup>j</sup> | ** | ** | -1.8E-02(4.3E-02)* | NS | 2.4(0.2)** | -3.1E-01(1.8E-01)** | 15.7(0.2)** | NA | NA |
| BS | ** | * | NS | NS | 4.1E-01(5.5E-02)** | -4.4E-01(5.3E-02)** | NA | NA | NA |
| Trait | Survivors |  |  |  |  |  |  |  |  |
|  | Sex | Tank | MA | MW | ADG0 | LogBL | BS | CG | HA |
| ADG0 | ** | ** | -2.1E-05(6.1E-06)** | 1.4E-04(1.3E-05)** | NA<sup>l</sup> | NA | NA | NA | NA |
| ADGi | ** | ** | -1.8E-05(2.5E-05)NS | -3.7E-05(6.4E-05)NS | 2.2E-01(1.1E-01)* | -3.0E-04(1.3E-04)* | NA | NA | NA |
| LogBL | NS | ** | 1.5E-03(7.8E-03)NS | 2.6E-02(2.3E-02)NS | -3.2(4.5)NS | NA | NA | NA | NA |
| Trait | Non-Survivors |  |  |  |  |  |  |  |  |
|  | Sex | Tank | MA | MW | ADG0 | LogBL | BS | CG | HA |
| ADG0 | NS | * | -2.6E-05(7.3E-06)** | 1.4E-04(1.8E-05)** | NA | NA | NA | NA | NA |
| ADGi | NS | NS | 7.1E-06(1.3E-05)NS | -9.7E-05(4.0E-5)** | -4.4E-01(7.5E-02)** | 1.2E-07(9.0E-08)NS | NA | NA | NA |
| LogBL | NS | NS | -3.4E-03(4.8E-03)NS | 1.4E-02(1.3E-02)NS | -5.0(2.7)NS | NA | NA | NA | NA |
| ST | ** | * | -1.5E-02(1.0E-01)NS | -7.9E-01(2.8E-01)NS | 5.6(0.5)** | -3.4E-01(3.9E-01)NS | NA | NA | NA |

± LogBL, ADG0, ADGi, ST and BS were not included into the models for HW as HW was recorded on relatives
<sup>a</sup> Marking age <sup>b</sup> Marking weight <sup>c</sup> Average daily gain prior infection <sup>d</sup> Logarithm of bacterial load <sup>e</sup> Binary survival <sup>f</sup> Contemporary group <sup>g</sup> Harvest age <sup>h</sup> Average daily gain during infection <sup>i</sup> Harvest weight <sup>j</sup> Survival time <sup>k</sup> Not significant <sup>l</sup> Not assessed \* p < 0.05 \*\* p < 0.001

### Effect of early growth and bacterial load on growth and survival traits under infection

As shown in table 2, ADG0 had a significantly positive effect (p < 0.001) on the traits ADGi, LogBL, ST and BS, implying that fast growth prior to infection also corresponded on average to faster growth and higher chance for survival during the infection process. This significance was also found in survivors (for ADGi, p < 0.05) and non-survivors (for ADGi and ST, p < 0.001) seperately. LogBL, in contrast, was found to have a significant negative effect on both survival traits BS and ST, and a negative, though not significant (p = 0.09) effect on ADGi (Table 2). The latter suggests that differences in growth during infection occur due to other factors than differences in bacterial load.

### Heritabilities and genetic correlations

Estimated heritabilities and genetic correlations obtained by including both survivors and non-survivors in the models, and by analyzing both categories separately, are presented in Table 3. A significant and moderate heritability was estimated for growth rate prior to the experimental challenge (h^2^ = 0.30 ± 0.05 in pooled dataset, and slightly lower values when survivors and non-survivors were analyzed separately (Table 3)). This estimate decreased considerably during the experimental challenge test (e.g. h^2^ = of 0.07 ± 0.02 in pooled dataset), which could be explained by a substantial increase in the phenotypic variance of ADGi compared to ADG0 (Table 3). Higher heritability estimates were obtained for HW of related individuals (0.38 ± 0.03 in pooled dataset). The survival traits ST and BS were found to be moderately heritable (0.16 ± 0.04 and 0.18 ± 0.03, for ST and BS respectively, in the full dataset, although heritability of ST dropped to 0.08 ± 0.04 when survivors were removed), when differences in ADG0 and LogBL were accounted for in the models. The estimate of heritability for LogBL was low and not significantly different from zero (0.04 ± 0.03) in the pooled dataset, but increased to 0.12 ± 0.05 and 0.11 ± 0.06, respectively, when survivors and non-survivors were analyzed separately.

**Table 3.** Genetic parameters and estimated heritabilities (SE), genetic and phenotypic correlations (below and above diagonal, respectively) for average daily gain prior infection (ADG0), during infection (ADGi), harvest weight (HW), bacterial load (logBL), day of death (ST) and binary survival (BS) for pooled, survivors and non-survivors animals

| Trait | Pooled animals |  |  |  |  |  |  |  |  |
| --- | --- | --- | --- | --- | --- | --- | --- | --- | --- |
|  | GenVar<sup>b</sup> | PhenVar<sup>c</sup> | ResVar<sup>d</sup> | ADG0 | ADGi | HW | Log BL | ST | BS |
| ADG0 | 0.08 ± 0.02 | 0.28 ± 0.01 | 0.20 ± 0.01 | **0.30(0.05)\*** | 0.05(0.02)* | 0 | -0.05(0.04) | 0.22(0.02)* | 0.10(0.02)* |
| ADGi | 0.11 ± 0.04 | 1.69 ± 0.05 | 1.58 ± 0.05 | 0.40(0.16)* | **0.07(0.02)\*** | 0 | -0.11(0.04)* | 0.12(0.02)* | 0.53(0.01)* |
| HW | 0.23 ± 0.02 | 0.59 ± 0.01 | 0.36 ± 0.01 | 0.64(0.09)* | -0.14(0.20) | **0.38(0.03)\*** | 0 | 0 | 0 |
| LogBL | 5.19E-03 ± 4.8E-03 | 0.14 ± 7.32E-03 | 0.13 ± 7.8E-03 | 0.09(0.43) | -0.23(0.50) | 0.63(0.28)* | **0.04(0.03)** | -0.07(0.04) | -0.50(0.03)* |
| ST | 14.3 ± 3.68 | 89.2 ± 2.89 | 74.9 ± 3.01 | 0.43(0.18)* | 0.15(0.28) | -0.50(0.13)* | -0.03(0.57) | **0.16(0.04)\*** | 0.70(0.01)* |
| BS | 0.22 ± 0.51E-01 | 1.23 ± 0.51E-01 | 1.00 ± 0.01 | 0.05(0.19) | 0.84(0.07)* | 0.23(0.16) | -0.97(0.10)* | 0.85(0.09)* | **0.18(0.03)\*** |
| Trait | Survivors |  |  |  |  |  |  |  |  |
|  | GenVar | PhenVar | ResVar | ADG0 | ADGi | HW | LogBL | ST | BS |
| ADG0 | 6.91E-08 ± 1.37E-08 | 2.57E-07 ± 1.31E-08 | 1.87E-07 ± 1.41E-08 | **0.27(0.05)\*** | 0.14(0.03)* | 0 | -0.01(0.05) | NA<sup>a</sup> | NA |
| ADGi | 2.72E-07 ± 1.38E-07 | 1.36E-06 ± 1.11E-07 | 1.10E-06 ± 1.15E-07 | 0.36(0.16)* | **0.20(0.09)\*** | 0 | -0.12(0.05)* | NA | NA |
| HW | 0.35 ± 2.05E-02 | 0.68 ± 0.11E-02 | 0.33 ± 0.11E-02 | 0.63(0.09)* | -0.16(0.19) | **0.52(0.02)\*** | 0 | NA | NA |
| LogBL | 1.24E-02 ± 5.75E-03 | 0.11 ± 4.27E-03 | 9.41E-02 ± 4.55E-03 | 0.39(0.60) | 0.11(0.77) | 0.25(0.32) | **0.12(0.05)\*** | NA | NA |
| ST | NA | NA | NA | NA | NA | NA | NA | **NA** | NA |
| BS | NA | NA | NA | NA | NA | NA | NA | NA | **NA** |
| Trait | Non-Survivors |  |  |  |  |  |  |  |  |
|  | GenVar | PhenVar | ResVar | ADG0 | ADGi | HW | LogBL | ST | BS |
| ADG0 | 5.82E-08 ± 1.97E-08 | 2.77E-07 ± 1.63E-08 | 2.19E-07 ± 1.80E-08 | **0.21(0.07)\*** | -0.27(0.03)* | 0 | -0.19(0.07)* | 0.33(0.03)* | NA |
| ADGi | 2.13E-08 ± 3.62E-08 | 1.46E-06 ± 6.58E-08 | 1.44E-06 ± 7.17E-08 | -0.50(0.57) | **0.02(0.03)** | 0 | -0.05(0.07) | 0.28(0.03)* | NA |
| HW | 0.35 ± 2.05E-02 | 0.68 ± 0.11E-02 | 0.33 ± 0.11E-02 | 0.58(0.13)* | -0.28(0.37) | **0.52(0.02)\*** | 0 | 0 | NA |
| LogBL | 3.39E-03 ± 2.04E-03 | 3.11E-02 ± 1.44E-03 | 2.77E-02 ± 1.59E-03 | 0.11(0.44) | -0.89(0.56) | -0.36(0.27) | **0.11(0.06)** | -0.11(0.06) | NA |
| ST | 5.98 ± 3.25 | 70.47 ± 3.27 | 64.48 ± 3.75 | 0.64(0.17)* | 0.54(0.54) | -0.35(0.20) | -0.09(0.41) | **0.08(0.04)** | NA |
| BS | NA | NA | NA | NA | NA | NA | NA | NA | **NA** |

\* p < 0.05
<sup>a</sup> Not assessed
<sup>b</sup> Genetic variance
<sup>c</sup> Phenotypic variance
<sup>d</sup> Residual variance

A significant and positive genetic correlation was estimated between ADG0 and ADGi in the full dataset (0.40 ± 0.16) and for survivors only (0.36 ± 0.16), whereas genetic correlations between these traits were not found significant for non-survivors. Estimates of genetic correlations between ADG0 and HW were favorable (0.64 ± 0.09 in pooled dataset) and robust across the datasets (Table 3). Growth prior to infection and ST also showed a significant and positive genetic correlation (0.43 ± 0.18 and 0.64 ± 0.17 in full dataset and non-survivors, respectively), indicating that fish with higher growth rates prior to infection are more likely to survive the infection. However, genetic correlations between ADG0 and BS were not found significantly different from zero (0.05 ± 0.19). Lastly, no significant genetic correlation was found between ADG0 and LogBL in either dataset.

Binary survival was strongly and favorably correlated with ST (0.85 ± 0.09), ADGi (0.84 ± 0.07) and LogBL (−0.97 ± 0.10), whilst no significant correlation with HW was found. Surprisingly, a significant unfavorable correlation was found between ST and HW (−0.50 ± 0.13), although the association became weaker and non-significant once the censored ST values of survivors were removed from the analyses (Table 3). Similarly, the statistically significant undesirable positive correlation between HW and LogBL estimated with the pooled dataset was no longer observed when the analysis was stratified into survivors and non-survivors (Table 3). Lastly, genetic correlations between ADGi and LogBL tended to be favorable, but were not found statistically significant in either of the datasets.

## Discussion

In order to include *P. salmonis* resistance into the breeding goal it is necessary to determine if the trait is heritable and its potential association with other economically important traits, such as growth. The current dataset comprises animals that were exposed to *P. salmonis* using an experimental challenge, and previously analyzed by Yáñez *et al.*, [12]. However, these authors only focused on the genetic association between harvest weight and day of death as a measure of resistance.

The current work provides novel insights into the genetic (co)variation between growth and *P. salmonis* resistance. By defining growth as average daily gain prior and during an infection with *P. salmonis*, we estimated heritabilities and its genetic correlation with resistance defined as survival time and binary survival, and the less commonly measured but epidemiologically important bacterial load. Moreover, estimates of the genetic correlation of all these traits with harvest weight was also determined using a large cohort from a coho salmon breeding population.

We found a moderate significant genetic variation for early growth rate (0.30 ± 0.05). Similar heritability values have been reported for growth rate in others salmonid species, ranging from 0.32 to 0.35 [35,36].

When growth rate was measured during infection with *P. salmonis*, heritability was up to six fold lower than the value for growth prior to infection. A similar drop in heritability for average daily gain during infection, compared growth rate prior infection have been observed in pigs [37] and chickens [38]. In our study, this drop in heritability could be explained by a relatively stronger increase in the phenotypic variance (with some fish losing rather than gaining weight due to infection), than in the genetic variance (Table 3). The results suggest that differences in growth under infection are primarily controlled by environmental rather than genetic factors, once individual differences in early growth or in disease resistance (represented by log-transformed bacterial load included as fixed covariate) are accounted for. Nevertheless, heritability estimates for growth under infection were still significantly different from zero, which is indicative for genetic variation in tolerance, in addition to resistance [25,31].

A moderate and positive genetic correlation was found between growth prior to and under infection. This favorable and significant genetic correlation was also estimated between growth prior to infection and harvest weight. The results indicate not only that fish with greater genetic growth potential at early stage in a pathogen-free environment in fresh water also tend to have greater growth potential during infection with *P. salmonis*, but also during the sea-rearing period, reaching higher body mass at harvest.

Significant additive genetic variation was estimated for ST and BS. These estimates are in agreement with previous estimates for the same and other types of pathogens for salmonid species (for a detailed review see [9]). Furthermore, a moderate and favorable genetic correlation between early growth and ST was found. These results corroborate findings indicating that fish with faster growth prior to and during infection are more likely to survive after an experimental challenge with a bacterial agent [39,40]. Hence, together the results of this study suggest that selection for early growth is expected to have a positive effect on growth under *P. salmonis* infection and harvest weight, without negatively impacting on survival.

One of the novelties of the present studies is the inclusion of bacterial load as additional measures of host resistance to infection. Even though pathogen load is commonly used as measure of disease resistance in domestic livestock [41–43], it is rarely used in aquaculture for practical reasons [44]. Measurements of individual pathogen load not only provide novel insights into different genetic response mechanisms to infection, such as resistance and tolerance or endurance [31,45,46] and their impact on survival [21], but may also help to predict potential epidemiological effects of selection, as individuals with high pathogen load may be more infectious.

In our study, the regression coefficient for logBL, when fitted into the statistical models for ST and BS, was significantly different from zero and negative, indicating that individuals with higher bacterial load were more likely to die and tended to die faster when infected with *P. salmonis*. Although the sample size for bacterial load in our study was too small to obtain accurate genetic parameter estimates, we found significant genetic variation for bacterial load in surviving animals (0.12 ± 0.05) and a similar borderline significant genetic variation in non-survivors (0.11 ± 0.06). Furthermore, a strong favorable genetic correlation was found between log-transformed bacterial load and binary survival, and genetic correlations between LogBL and ST or growth traits tended to be negative, suggesting that selection for growth or survival post *P. salmonis* infection will not simultaneously increase pathogen load.

In our study, final body weight used to calculate ADGi, and BL were measured at time of death for non-survivors and at the end of the trial for survivors. This implies that the trait measurements may relate to different stages of infection in survivors and non-survivors. The low genetic and phenotypic correlations for these traits measured in survivors and non-survivors indicate that these traits should be considered as biologically different in both groups of individuals. Indeed, survivors may have already fully or partially recovered from infection at the time of recording and may thus have had reduced bacterial load in contrast to non-survivors whose bacterial load may have peaked at the time of death. Similarly, in the case of ADGi, non-survivors fish may have died when body weight reached a minimum, whereas survivors may have experienced compensatory growth at the later stages of the experiment. For these reasons, the analyses were carried out with data from survivors and non-survivors pooled (with BS fitted as fixed effect to partly account for these differences) in order to maximize statistical power, and for survivors and non-survivors separately to disentangle the effects of confounding with recording times. Furthermore, genetic parameter estimates may be slightly upward biased due to the fact that bacterial load was only measured in families from the extreme ends of the survival time breeding value distributions.

Nevertheless, results of the genetic estimates within survivors and non-survivors analyzed separately were overall consistent with those obtained with pooled animals, although standard errors were higher. In particular, growth prior to infection showed a generally favorable correlation with growth and survival during *P. salmonis* infection, and harvest weight, and no robust antagonistic genetic relationship between growth, survival and bacterial load was identified in this study.

From a resource allocation theory point of view, a negative correlation between resistance and growth would be expected, given that these are two competing resource-demanding mechanisms [47]. Indeed, previous studies found a negative genetic correlation between body weight and resistance (as day of death) to SRS and *viral haemorrhagic septicaemia* (VHS) in salmonid species [12,15,48]. Our current results do not support this trade-off, as neither ADG0 or ADGi were antagonistically related to survival. Instead, the estimated positive and favorable genetic correlations in pooled and non-survivors individuals, suggest that fish with higher genetic growth rate measured in freshwater at early stage are also genetically more resistant to *P. salmonis*. Similar results have been obtained in Atlantic salmon and rainbow trout [39,49]. This trade-off was only observed due to the unfavorable genetic correlation between ST and HW when the former was measured in pooled animals. However, the genetic correlation between ST and HW was not significantly different from zero when only non-survivors animals were used, suggesting a less robust estimation compared to ST and ADG0, which was positive and significantly different from zero when using only susceptible animals. Furthermore, we found a favorable genetic correlation for ADG0 with respect to ST and HW, which indicate a positive relationship between early growth in fresh water (ADG0), late growth in seawater (HW) and survival time (ST).

Differences at the development of the immune system at early life stages, given by body size at time of infection may explain the lack of trade-off [39]. Furthermore, the role of insulin-like growth factor (IGF) could play a key role as has been associated with increased survival and detected in higher levels in faster growing fish [50–52].

Previously, an up-regulation of pro-inflammatory genes has been detected in Atlantic salmon families with early mortality following a *P. salmonis* infection [16,30,53]. Moreover, using genome-wide association studies, candidate genes related with pro-inflammatory response proximate to markers associated with *P. salmonis* survival were identified in Atlantic [16] and Coho salmon [13]. In our work, fish from the 17 more susceptible families experienced a higher weight loss than those from the 16 most resistant families, (p < 0.0001, data not shown). We propose that an exacerbated, ineffective inflammatory response may have led to tissue damage and the subsequent weight reduction in these individuals, with subsequent mortality. However, further studies are necessary to test these relationships between immune response and weight lost in coho salmon. The availability of a coho salmon reference genome (assembly accession = GCA_002021735.1) and the international initiative on Functional Annotation of All Salmonid Genomes (FAASG), will facilitated further functional studies in salmonid species [54].

One potential limitation of the current study refers to the censored data. The current censored distribution for mortality violates the normality assumption for the linear mixed models. Moreover, ignoring the censoring could cause slight bias in the estimates. Repeating the analyses for uncensored non-survivors only could partly overcome this problem, but at the loss of statistical power. Alternatively, this situation can be partly overcome with using survival analysis (e.g. proportional hazard frailty models) [6]. However, bivariate analyses with such models present some difficulties in terms of fitting and interpretation and therefore linear mixed models are often considered more robust in this case. From the genetic improvement perspective, predictive ability can be considered more relevant than goodness of fit for a given model. In this regard, it has been found that when comparing proportional hazard frailty models with linear mixed models, the increase in accuracy of selection is marginal in the case of *P. salmonis* resistance in Atlantic salmon [14].

Finally, although survival to infection is generally considered as desirable breeding goal in aquaculture, from an epidemiological point of view, fish that survive an infection, but that harbor and shed a large amount of infectious pathogens may be highly undesirable as they are more likely to infect others. The results of this study would suggest that selection for survival to *P. salmonis* infection will not simultaneously increase bacterial load. However, recent infection studies in Turbot demonstrated that survival is a composite genetic trait influenced by genetic variation in host resistance, tolerance and infectivity [27,55]. Future studies that provide a deeper understanding of the underlying mechanisms and their genetic regulation affecting survival of coho salmon to *P. salmonis* infection, are therefore warranted.

## Conclusion

The current study showed the presence of significant genetic variation for average daily gain in an early stage of a coho salmon life cycle. This genetic variation decreased during infection by the facultative intracellular bacteria *Piscirickettsia salmonis*, and a moderate positive genetic correlation between growth prior and during infection was observed. We identified that early growth is positive genetically correlated with *P. salmonis* resistance measured as day of death and with harvest weight. Furthermore, we found no robust antagonistic genetic relationship between growth, survival and bacterial load.These results suggest that selective breeding for early growth, can indirectly improve harvest weight and resistance to *P. salmonis* in the current population. To our knowledge this is the first study elucidating significant genetic variation for pathogen load in salmonid species as a measurement of resistance, and its genetic correlation with commercially important traits.

## Declarations

### Ethics approval and consent to participate

Experimental challenge and sample procedures were approved by the Comité de Bioética Animal from the Facultad de Ciencias Veterinarias y Pecuarias, Universidad de Chile (Certificate N08-2015).

### Consent for publication

Not applicable

### Availability of data and materials

The dataset used during the current study is commercially sensitive and could be available from the corresponding author on reasonable request.

### Author’s contribution

JY and JL conceived the experiment and provided data for analysis. AB performed the analysis. AW and RH helped to optimize the analysis. AB, AW, RH, and JY interpreted the results. AB and AW drafted the manuscript. All authors improved the writing, read and approved the final manuscript.

## Acknowledgements

AB want to acknowledge to the National Commission of Scientific and Technologic Research (CONICYT) for the funding through the National PhD funding program.

